# Using genomic scars to select immunotherapy beneficiaries in advanced non-small cell lung cancer

**DOI:** 10.1101/2022.09.23.509152

**Authors:** H. C. Donker, B. van Es, M. Tamminga, G. A. Lunter, L. C. L. T. van Kempen, E. Schuuring, T. J. N. Hiltermann, H. J. M. Groen

## Abstract

In advanced non-small cell lung cancer (NSCLC), response to immunotherapy is difficult to predict from pre-treatment information. Given the toxicity of immunotherapy and its financial burden on the healthcare system, we set out to identify patients for whom treatment is effective. To this end, we used mutational signatures from DNA mutations in pre-treatment tissue. Single base substitutions, doublet base substitutions, indels, and copy number alteration signatures were analysed in *m* = 101 patients (the discovery set). We found that tobacco smoking signature (SBS4) and thiopurine chemotherapy exposure-associated signature (SBS87) were linked to durable benefit. Combining both signatures in a machine learning model separated patients with a progression-free survival hazard ratio of 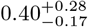 on the cross-validated discovery set and 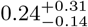 on an independent external validation set (*m* = 56). This paper demonstrates that the fingerprints of mutagenesis, codified through mutational signatures, select advanced NSCLC patients who may benefit from immunotherapy, thus potentially reducing unnecessary patient burden.

## Introduction

In non-small cell lung cancer (NSCLC), response to immunotherapy is low. Radiology-assessed response is typically around 20-25% [1], while the percentage of patients achieving durable benefit (DB), defined as progression-free survival (PFS) ≥ 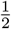 year, is only slightly higher. Predictors can help to narrow down specific subpopulations for which treatment is particularly effective. Programmed death ligand 1 (PD-L1) protein expression in tumor tissue, and tumor mutational burden (TMB), defined as the number of acquired amino acid sequence-changing mutations [2], is currently used to predict the efficacy of immunotherapy in NSCLC [3]. The explanation behind the predictive value of TMB is that the accumulation of mutations in coding DNA results in a high diversity of neoantigens. In turn, these neoantigens induce a broad anti-tumor adaptive immune response. *In silico* analysis of neoantigens found a near-perfect relation between TMB and inferred number of neoantigens [4]. Once the cancer cell expresses neoantigens, the cancer cell can be eliminated through immune recognition and cell killing [1, 2].

While several studies show a clear association of TMB with response to treatment, not all do [5], motivating the search for an improved proxy for treatment efficacy. The chief advantage of TMB is that it is easy to compute by pooling all amino acid sequence-changing mutations from whole-exome sequencing data [2], irrespective of their genomic context. However, this ignores the fact that somatic mutations are generated by a range of processes. Nucleotide context is a clue to the genesis of mutations [6]. Analysis of mutation spectra, which partition mutations by alteration and DNA context, have revealed that both exogenous and endogenous DNA mutational processes can be linked to specific signatures [7, 8, 9]. Over the last decade, increasingly large datasets, such as the COSMIC database, have enabled the systematic identification of mutational signatures [10, 11]. And in many cases, also the elucidation of their aetiology, particularly for single base substitutions [7]. For instance, specific DNA mutational signatures or “genomic scars” left by e.g. polycyclic aromatic hydrocarbons [12], ultraviolet light exposure [13], and platinum chemotherapy [14] have all been validated in experiments.

Evidence is accumulating that mutational signatures in cancer are therapeutically relevant [9]. Given the partial success of TMB for predicting immunotherapy efficacy in NSCLC [2], a logical next step is to try and improve TMB by sieving out irrelevant mutations. Mutational signatures, which disentangle mutations by their proposed root cause, may help to pinpoint relevant alterations. In fact, specific mutational signatures (e.g., APOBEC A3A) have been hypothesised to be promising candidates for immune stimulation treatments [15]. Along this line, previous studies performed *de novo* identification of single base substitution signatures in NSCLC and linked these to immunotherapy efficacy [16, 17]. Given the relatively small datasets used in these studies, only three signatures were identified in both cases [16, 17]. Specifically, Wang *et al*. [16] found that patients with durable clinical benefit were enriched in signatures with similarities to COSMIC signatures SBS2 and SBS13 that are associated with damage from APOBEC, a family of enzymes that are part of the innate anti-retroviral defense that operates by generating mutations in single-stranded DNA [18, 9]. Signatures similar to clock-like singlet SBS1, capturing mutations that steadily accrue with age, were linked to non-response to immunotherapy by Chong *et al*. [17].

Here, instead of *de novo* analysis we directly use previously catalogued mutational signatures like in earlier work [4, 19]. We expand previous efforts by interrogating an order of magnitude more signatures, including the recently developed copy number signatures [20, 21], for their role in eliciting an immune response. In addition, we validate the presence of single base substitution signatures at the RNA level. The primary goal of this work is to develop and validate an immunotherapy efficacy model for advanced NSCLC patients to help reduce ineffective treatment. We hypothesise that the genomic scars in pre-treatment tumor tissue, decomposed into mutational signatures, can help identify durable immunotherapy beneficiaries (Fig. 1a).

**Figure 1:**
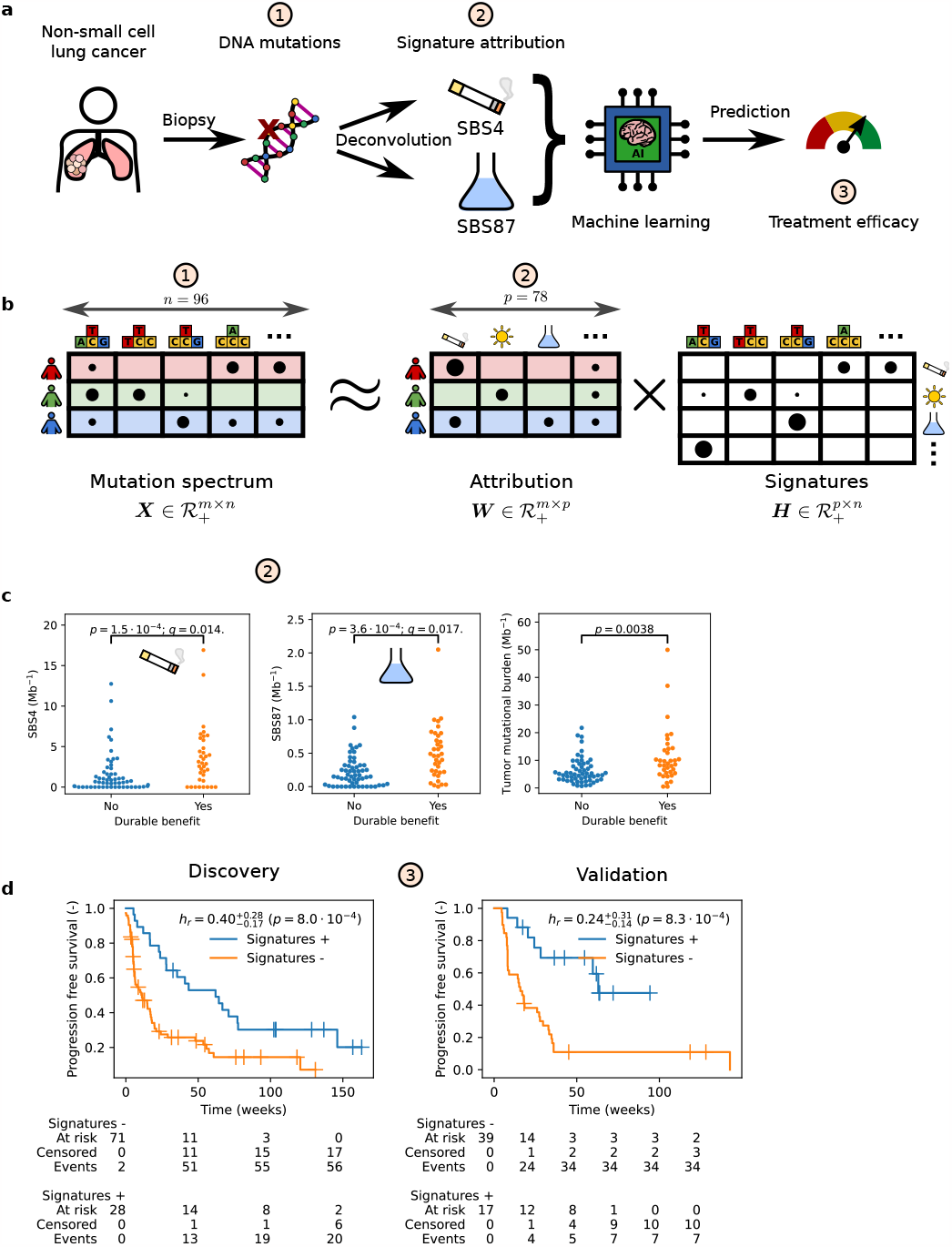
Mutational signatures from pre-treatment tumor tissue predict immunotherapy efficacy in advanced non-small cell lung cancer. **a**, Single base substitutions (SBS) are determined from preimmunotherapy tumor material. After deconvolution into signature attributions, a machine learning classifier uses smoking-associated signature SBS4 and thiopurine chemotherapy-associated signature SBS87 to predict durable benefit (DB) from immunotherapy. **b**, Cartoon illustration of SBS signature deconvolution, where we solve for signature attribution ***W*** given mutation spectrum ***X*** and COSMIC signatures ***H*** through ***X*** ≈ ***W H***. Nucleotide pyramids indicate SBS with flanking context; Sun, cigarette, and Erlenmeyer symbols depict example aetiologies; Shading highlights information that pertains to the corresponding patient. For illustration purposes, the size of the dots do not represent actual data. **c**, Signatures SBS4 [*q* = 0.014, Benjamini-Hochberg corrected Kolmogorov-Smirnov (B-HK-S) test] and SBS87 (*q* = 0.017, B-HK-S test) are linked to DB (discovery set). For reference, tumor mutational burden is also shown. **d**, Patients predicted to have DB (Signatures +, blue line) have superior progression-free survival compared to those predicted to have non-DB (Signatures -, orange line) in the discovery set (left panel). The classifier’s performance replicates in an independent validation set (right). Censored observations are indicated by crosses. Estimates and corresponding 95% confidence intervals are indicated by sub and superscripts.

## Result & Discussion

In total, *M* = 157 patients with advanced NSCLC were analysed, *m* = 101 in the discovery and *m* = 56 in a separate validation set. The population had a mean age of 63.2, consisted of an approximately equal split between males and females (53.5%). The majority of the patients (85/147, 57.8%) did not achieve durable benefit from immunotherapy (Table 1). In the Discovery dataset, 21 patients (20.8 %) received combination therapy.

**Table 1:** Patient characteristics and outcome of advanced non-small cell lung cancer. Symbols and abbreviations: *α*CTLA-4, cytotoxic T lymphocyte-associated antigen-4 inhibitor; *α*PD-1, programmed death-1 inhibitor; *α*PD-L1, programmed death-ligand 1 inhibitor; *α*TNFRSF7, tumor necrosis factor receptor superfamily type 7 inhibitor; *α*VEGF-A, vascular endothelial growth factor inhibitor; *μ*, mean; *σ*, standard deviation; DR, data request; *Q*_1_, first quartile; *Q*_3_, third quartile.

|  |  | Discovery (Hartwig DR #094) | Validation [Miao *et al.* ('18)] | Overall |
| --- | --- | --- | --- | --- |
| $m$ | | 101 | 56 | 157 |
| Age, $\mu$ ($\sigma$) | | 63.84 (8.72) | 61.49 (8.75)* | 63.16 (8.77) |
| Gender, $m$ (%) | female | 52 (51.49) | 32 (57.14) | 84 (53.50) |
|  | male | 49 (48.51) | 24 (42.86) | 73 (46.50) |
| Smoker, $m$ (%) | current | | 14 (25.00) | 14 (8.92) |
|  | former |  | 29 (51.79) | 29 (18.47) |
|  | never |  | 13 (23.21) | 13 (8.28) |
|  | unknown | 101 (100.00) |  | 101 (64.33) |
| Prior therapy, $m$ (%) | chemotherapy | 33 (32.67) | | 33 (21.02) |
|  | naive | 18 (17.82) |  | 18 (11.46) |
|  | radiotherapy | 7 (6.93) |  | 7 (4.46) |
|  | radiotherapy + chemotherapy | 43 (42.57) |  | 43 (27.39) |
|  | unknown |  | 56 (100.00) | 56 (35.67) |
| Treatment, $m$ (%) | αCTLA-4 + αPD-1 | 1 (0.99) | | 1 (0.64) |
| | αCTLA-4 + αPD-1 with chemotherapy | 1 (0.99) | | 1 (0.64) |
| | αPD-1 | 77 (76.24) | | 77 (49.04) |
| | αPD-1 with chemotherapy | 13 (12.87) | | 13 (8.28) |
| | αPD-1/αPD-L1 | | 56 (100.00) | 56 (35.67) |
| | αPD-L1 | 3 (2.97) | | 3 (1.91) |
| | αPD-L1 with chemotherapy | 2 (1.98) | | 2 (1.27) |
| | αPD-L1/cabozantinib | 1 (0.99) | | 1 (0.64) |
| | αTNFRSF7 + αPD-1 | 1 (0.99) | | 1 (0.64) |
| | αVEGF-A + αPD-1 | 2 (1.98) | | 2 (1.27) |
| Durable benefit, $m$ (%) | no | 57 (56.44) | 28 (50.00) | 85 (54.14) |
|  | yes | 36 (35.64) | 26 (46.43) | 62 (39.49) |
|  | unknown | 8 (7.92) | 2 (3.57) | 10 (6.37) |
| Sequencing, $m$ (%) | whole exome | | 56 (100.00) | 56 (35.67) |
|  | whole genome | 101 (100.00) |  | 101 (64.33) |
| Tissue, $m$ (%) | formalin-fixed paraffin-embedded | | 56 (100.00) | 56 (35.67) |
|  | fresh frozen | 101 (100.00) |  | 101 (64.33) |
| Tumor purity, median [$Q_1, Q_3$] | | 0.41 [0.28, 0.56] | 0.35 [0.24, 0.54] | 0.39 [0.27, 0.56] |

\* Age was missing for 15 patients in the validation cohort.

### Mutational signatures in the discovery set

Whole genome sequencing revealed a mean of 7.80 Mb^−1^ [median 5.43 Mb^−1^; inter quantile range (IQR) 3.90-9.85 Mb^−1^] non-synonymous variants in the discovery set, consistent with previous reports on whole exome sequencing [22, 23].

Mutational signature data of *p* = 47 single base substitutions (SBS), *p* = 11 doublet base substitution (DBS), *p* = 17 short insertion and deletion (indel), and *p* = 20 copy number signatures were determined (see Fig. 1b for a schematic overview) and analysed, after discarding sequencing artefact attributed signatures (Supplementary Table 1). Ranking mutational signature attribution ***W*** (see the Methods for details) by the median (across samples) shows that nine SBS signatures are present in the majority of samples (Fig. S1a, Supplementary Material), with signature SBS4, a lung cancer-specific signature [24] that is linked to tobacco smoking [6, 12, 7], having the highest median value (0.91 Mb^−1^), although exhibiting considerable variability (IQR: 0.00-3.14 Mb^−1^). Signature attributions of indel and doublet base substitution were an order of magnitude lower (Fig. S1b, Supplementary Material) with the two highest median signatures, ID3 [7, 10] (median: 0.11 Mb^−1^, mean: 0.17 Mb^−1^) and DBS2 [25, 7] (median: 0.06 Mb^−1^, mean: 0.10 Mb^−1^), both attributed to tobacco smoking, like SBS4. Whole genome copy number deconvolution revealed that homologous recombination deficiency [21] signature CN17 had the highest median attribution (median: 37.2, mean: 46.5, Fig. S1c, Supplementary Material).

### Mutational signatures linked to durable benefit

Next, we looked for pre-immunotherapy mutational signatures that were determinants of therapy efficacy. Univariate analysis (*m* = 93 patients) singled out two single base substitution signatures (Fig. 1c). Tobacco smoking signature SBS4 was significantly different (*q* = 0.014, B-HK-S test) in patients who derive DB from immunotherapy. This finding underpins earlier work that found that smoking-attributed transversion-high tumors [26], or enrichment in smoking signature [19], had improved outcome in ICI-treated NSCLC. Similarly, SBS87— whose mutations coincide with thiopurine chemotherapy exposure [27, 9]—was also found to differ between both groups of patients (*q* = 0.017, B-HK-S test). Note that, according to the clinical records available to us, none of the patients has been treated with thiopurine-related compounds.

We subsequently considered the mutual dependence between SBS4 and SBS87. Part of this correlation is mediated through the outcome (DB versus non-DB) and the total number of amino-acid sequence-changing mutations (i.e., TMB). We, therefore, normalised both signatures by TMB and correlated separately for patients with DB and non-DB. In both outcome groups, signatures SBS4 and SBS87 were unrelated (Kendall *τ* = − 3.6 · 10^−2^, *p* = 0.76 and *τ* = − 8.2 · 10^−3^, *p* = 0.95 for DB and non-DB, respectively).

Earlier work linked clock-like mutational signature SBS1—capturing substitutions that steadily accrue with age—with non-response and worse survival [17]; we could not replicate the association with DB, even without multiple testing correction (*p* = 0.36, K-S test). Another study linked mutational signatures associated with APOBEC—a family of enzymes that are part of the innate anti-retroviral defense that operates by generating mutations in singlestranded DNA [18, 9]—with improved immunotherapy outcome [16]. Validation of APOBEC mutational signatures SBS2 and SBS13 [16] showed a trend for SBS13 (*p* = 0.04 K-S test, *q* = 0.76 B-HK-S test) but not for SBS2 (*p* = 0.66 K-S test, *q* = 1.0 B-HK-S test). Given that APOBEC mutagenesis is highly transient, with episodic bursts of mutations [28], our samples, which represent a snapshot in time, are perhaps less suited to fully interrogate the relevance of this mutational signature on treatment outcome. None of the doublet, indel, and genome-wide copy number alteration signatures was significantly associated with DB.

### A signature-based classifier predicts immunotherapy benefit

In combining SBS4 and SBS87 signature attributions (representing, per signature, the number of amino-acid sequence changing singlets in DNA) both remained significant in each cross-validated fold, confirming that the aforementioned univariate analysis does not lead to overfitting on the discovery set. The classifier scored an area under the receiver operating characteristic curve (ROC AUC) of 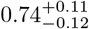 on the hold-out folds (Fig. S3, Supplementary Material). This was significantly higher than when the classifier was trained on TMB (ROC AUC: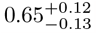, *p* = 0.016 PPT, Fig. S3a, Supplementary Material). The ROC curves of individual signatures were similar to that of the model (Fig. S4). With an estimated 43.8% patients with DB [29] as our classification probability threshold, a sensitivity of 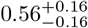, a specificity of 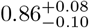, and an accuracy of 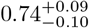was achieved (Table 2) with 8 false positive and 16 false negative classifications (Supplementary Table 3). The classifier was not perfectly calibrated (Fig. S5a), as expected from the conditional independence assumption of the naive Bayes model.

**Table 2:** Durable benefit (DB) classifier performance metrics and *p*-value comparing the performance between the two datasets. Abbreviations: ROC AUC, area under the receiver operating characteristic curve; AP, average precision; *h*_*r*_, hazard ratio of progression-free survival (PFS) comparing patients predicted DB versus predicted non-DB. Estimates and corresponding 95% confidence intervals are indicated by sub and superscripts.

| | ROC AUC | AP | $F_1$ | Sensitivity | Specificity | Accuracy | $h_r^*$ |
| --- | --- | --- | --- | --- | --- | --- | --- |
| **Discovery** |  |  |  |  |  |  |  |
| (Hartwig DR #094) | $0.74^{+0.11}_{-0.10}$ | $0.63^{+0.18}_{-0.15}$ | $0.63^{+0.14}_{-0.16}$ | $0.56^{+0.16}_{-0.17}$ | $0.86^{+0.09}_{-0.10}$ | $0.74^{+0.09}_{-0.10}$ | $0.40^{+0.28}_{-0.17}$ |
| **Validation** |  |  |  |  |  |  |  |
| (Miao *et al.* ('18)) | $0.69^{+0.13}_{-0.15}$ | $0.71^{+0.16}_{-0.19}$ | $0.57^{+0.17}_{-0.19}$ | $0.46^{+0.21}_{-0.21}$ | $0.86^{+0.11}_{-0.14}$ | $0.67^{+0.13}_{-0.13}$ | $0.24^{+0.31}_{-0.14}$ |
| $p$ | 0.57 | 0.56 | 0.68 | 0.54 | 1.00 | 0.35 | 0.32 |

\* All patients were included in the PFS analysis. For all other classification metrics, only patients where censoring permitted unambiguous DB label assignment were analysed.

To incorporate patients censored prior to the 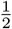 year mark, we compared predicted versus actual outcome using Kaplan-Meier (Fig. 1d) in all *m* = 101 discovery patients. Durable benefit predicted patients had superior median progression-free survival (62 for predicted DB versus 11 weeks for predicted non-DB). Cox regression provided additional confirmation that the classifier significantly predicted outcome based on mutational signatures SBS4 and SBS87 combined (hazard ratio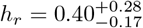, *p* = 8.0 · 10^−4^).

Independent validation on the data from Miao *et al*. [4] reproduces the classifier’s performance. The median survival in the external validation set (*m* = 56 patients) was 63 versus 16 weeks for DB-predicted patients compared to the others (Fig. 1d). All performance metrics were similar (Table 2), highlighting the reproducibility of our approach despite differences in (i) tissue handling (formalin-fixed paraffin-embedded versus fresh frozen), (ii) chemistry (whole exome versus genome capture), and (iii) bioinformatics pipeline. Unlike the discovery set, the ROC AUC of a model trained on TMB was not significantly different from the mutational signature model (ROC AUC 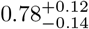 versus 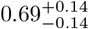, respectively, *p* = 0.18 PPT, Fig. S3b, Supplementary Material). This discrepancy was attributed to a difference in TMB distribution between the discovery and validation set (*p* = 0.031, KS test), while both SBS4 and SBS87 signature attributions remained similar in both cohorts (*p* = 0.082 and *p* = 0.69, respectively, KS test). A fair head-to-head comparison between TMB and the mutational signature approach requires a separate, substantially larger, study with more detailed patient characteristics and medical history than presented here (together with a more harmonised tissue handling, sequencing chemistry, and upstream bioinformatics preprocessing).

Error analysis of the discovery set revealed that treatment-naive patients were overrepresented in the top ten worst predicted false negatives (*p* = 0.015, Fisher exact test). Exclusion of (*m* = 18) treatment-naive patients slightly improved the model 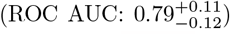on the discovery set. More strikingly, after training on pre-treated patients only, generalization on the validation patients—of whom prior therapy was unknown—also improved 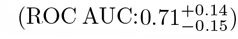. Caution is therefore warranted when applying the classifier to an (in our study, underrepresented) treatment-naive population.

Subanalysis of patients with smoking status (*m* = 54, validation set only, Table 1) showed that never smokers were more difficult to classify (ROC AUC: 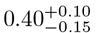 versus 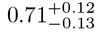, p = 0.048 UPT), although the numbers were low with only *m* = 3 durable beneficiaries in the never-smokers group.

Note that current/former smokers are known to have a better overall response rate [30]. Seeing that the performance was lower on the combined set indicates that the mutational signatures provide information that is orthogonal (or, complementary) to smoking status.

### Decision curve analysis

Net benefit is a decision-theoretic concept that quantifies the practical utility of a classifier [31, 32]. The integrated net benefit of using the signature-based model is positive (Figs. S6 and S7, Supplementary Material), with a median integrated combined net benefit (see e.g. Talluri *et al*. [33]) of 0.37 (IQR: 0.366-0.379) and 0.29 (IQR: 0.284-0.296) with respect to net benefits less than zero when treating *all* patients for the discovery and validation datasets, respectively. In general, according to Ref. [32] a model can be recommended for clinical use if, across a range of clinically reasonable probability thresholds, it has the highest level of benefit. This range of thresholds can be viewed as the acceptable range of *number-needed-to-treat* (NNT) to have one effective treatment and is clinician/protocol dependent. In Figs. S7 and S8, we see a net benefit for both the treated/untreated with respect to treating all patients for a probability threshold range of roughly 0.3-0.6 and 0.4-0.6 for the discovery and validation datasets respectively. Important to note, although we have fairly poor calibration for the 0.3-0.6 range within which we expect a relative net benefit with respect to the baseline, outside this range the net benefit is roughly equal. That is, within a probability threshold range of 0.3-0.6, so an NNT-range of roughly 1.5-3 pa-tients, we have a net benefit when applying our model. Outside this range, there is no added benefit.

### Relation of signature specific mutations to specific genomic loci

To better understand why SBS4 and SBS87 relate to DB, we set out to link their attribution to specific genes (in the Discovery set, *m* = 101 patients). Neither the SBS4 nor the SBS87 signature was correlated to the number of mutations in any of the 23 significantly lung cancer mutated genes in TCGA [22, 23] (Supplementary Table 2). Expanding the search from 23 to the top 2.5%, 5% and 10% highly expressed genes also found no correlation. In contrast, analysis of mutations in 523 genes contained in the clinically relevant TSO500 panel, consisting of genes canonically mutated in cancer, yielded eight genes [*ATM* (*q* = 0.029), *EPHA5* (*q* = 0.013), *LRP1* (*q* = 5.3 · 10^−4^), *MTOR* (*q* = 0.034), *NRG1* (*q* = 0.036), *PTPRD* (*q* = 1.7 · 10^−5^), *PTPRT* (*q* = 6.7 · 10^−5^), *RUNX1T1* (*q* = 0.023), Kendall *τ* correlation test] in which mutation count correlated significantly with smoking signature SBS4 (Fig. 2) after multiple testing correction and exclusion of non-mutated genes. When combined, the aggregated mutation count was also directly linked to DB (*p* = 0.0050, K-S test) in addition to the indirect correlation through SBS4. Four of these genes [namely, *LRP1* (*q* = 0.021), *PTPRD* (*q* = 0.021), *PTPRT* (*q* = 0.0091), and *RUNX1T1* (*q* = 0.039), but no other genes] also correlated with SBS87. Overall, significant genes were large, ranging from 146 Kb (ATM) to 2.3 Mb (PTPRD) [34, 35]. This was expected since in order to (significantly) correlate, enough mutations must be detected (the correlation with only zeroes is trivially zero). And the larger the gene, the more mutations can accumulate randomly. Functionally, both *ATM* and *EPHA5* interact at the site of DNA repair. Adding ATM to a DB logistic regression model with TMB changed the regression coefficient by more than 10 % (0.156 versus 0.117) indicating that ATM (but not EPHA5) potential confounds TMB. *MTOR* (a paralog of ATM) regulates cellular metabolism and the others are tumor suppressor genes. They are involved in cell interactions such as the *PTPRT* and suppression of inflammatory responses such as *STAT3. PTPRD* show deleterious mutations in 9% of lung cancers. *NRG1* suppresses the transcription of inflammatory cytokines and was the only gene (out of all eight) that was significant (*p* = 0.020) when added to a logistic regression model with TMB. Compared to TMB, mutations in NRG1 anti-correlated with DB (as indicated by the negative regression coefficient -2.6). One explanation could be that mutations in NRG1 disrupt the adaptive immune system’s capacity to induce an immune response. SBS87 only correlated with tumor suppressor genes. However, statistically, no enrichment for tumor suppressor or oncogenes was found in the set correlating with SBS4 (*p* = 0.25 and *p* = 0.16, respectively, Fisher exact test) nor with SBS87 (*p* = 0.08 and *p* = 0.16, respectively, Fisher exact test) relative to the TSO500 gene set.

**Figure 2:**
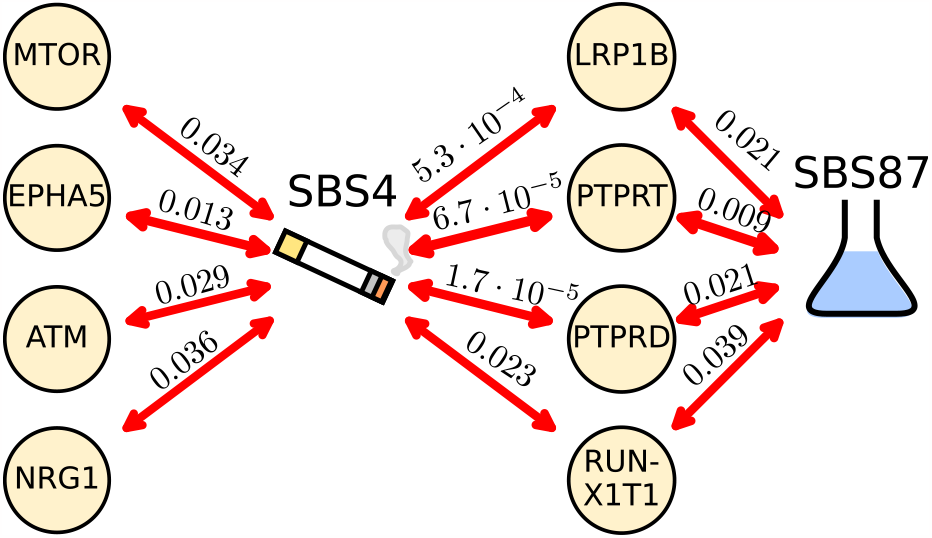
Signatures SBS4 and SBS87 correlate with mutations in genes canonically mutated in cancer (discovery set). Correlations were assessed using a Benjamini-Hochberg corrected Kendall *τ* (corrected p-values along the arrows) and the correlation strengths are indicated by the arrow line widths.

### Expression of SBS4/SBS87 mutations is not a sufficient condition for predicting durable benefit

Next, we aimed to explain the predictiveness of SBS4 and SBS87 attributed mutations by looking for differences at the RNA level. Assume that immune recognition, through the presentation of mutated protein fragments on the major histocompatability complex, is a prerequisite for eliciting an immune response. If transcription is a necessary condition for immune recognition of mutated DNA then, by extension, it must also be a necessary condition for predicting DB, assuming immunotherapy operates through an adaptive immune response. Focussing on transcripts with a SBS4 and/or SBS87 dominant variant (a signature-dominant variant accounts for ≥ 50% of the signature’s mutations, see Sec. A.4, Supplementary Material), we asked whether their transcription is also a sufficient condition for predicting DB.

We, therefore, did a subanalysis of paired RNA and DNA samples (*m* = 36 patients, Discovery dataset). To ensure that the subanalysis does not introduce a bias in the statistical analysis, we first verified that durable benefit (*p* = 0.67, Fisher exact test) and both SBS4 and SBS87 signatures (*p* = 0.30, *p* = 0.28 respectively, K-S test) did not differ from the entire discovery population. Next, comparing the number of transcripts containing an SBS4 or SBS87-dominant singlet versus transcripts containing any other singlet shows that the RNA abundance is similar in both groups (*p* = 0.71, K-S test, Fig. S2a, Supplementary Material). Compared to DB, the distribution of transcripts harbouring a smoking-associated signature SBS4-dominant variant was different from patients with non-DB (*p* = 0.0036, K-S, see Fig. S2b, Supplementary Material). However, this difference could be attributed to the number of SBS4 DNA mutations (Fig. S2c, Supplementary Material). For the thiopurine chemother-apy associated SBS87 signature, no difference in absolute number (*p* = 0.11, K-S, Fig. S2d, Supplementary Material) nor relative to the number of corresponding variants (*p* = 0.48, K-S, Fig. S2e, Supplementary Material) was found in the number of mutated transcripts between patients with and without DB. Differential gene expression of the mutated RNA of the aforementioned eight significant genes (*m* = 22 patients with paired DNA, RNA, and ≥ 1 mutations in any of these eight genes) detected no difference in the amount of mutated RNA between patients with and without DB (*q >* 0.05 for all genes, B-HK-S test). Together, these results show no evidence that distinguishes SBS4 and SBS87 from other mutations at the transcription level in durable beneficiaries. While these negative findings could point to the differences (i) being manifested further downstream immune processes or (ii) the capacity to bind to a broader spectrum of HLA alleles [36], we do not rule out that we lacked insufficient variant coverage at the RNA level (which was on average 19.8× for all and 19.1× for SBS4/SBS87 dominant variants) to detect subtle differences.

## Conclusion

We have identified tobacco smoking signature SBS4 and the recently identified thiopurine chemotherapy exposure-associated signature SBS87 [27, 9] as factors that are predictive of benefit from immunotherapy. Both signatures are linked to mutations in genes involved in DNA repair and/or the function of tumor suppressor genes that have a role in cellular immunological interactions and inflammatory cytokines. In contrast, none of the doublet base substitution, indel, or copy number alteration signatures were associated with durable benefit from immunotherapy. RNA analysis of the two signatures found no evidence that distinguishes these from other mutations in terms of immunotherapy efficacy.

When combined, these two signatures can help to select advanced NSCLC patients who may benefit from (combination) immunotherapy using information available prior to treatment initiation. These signatures were not tested in the context of other therapeutic interventions. As such, further investigations are required to validate if this signature is therapy-specific or if it may have prognostic value. The advantage of our predictor is that it inherits the mechanistic grounding of mutational signatures. However, more research is needed to establish if our approach reliably generalises to the (underrepresented) treatment-naive and non-smoking patient populations, or if additional adjustments are needed.

## Online Methods

### Cohort assembly

We retrospectively compiled clinical records and data of tumor and matched normal tissue (collected prior to treatment initiation) of immunotherapy-treated advanced NSCLC patients. To this end, patient characteristics, whole genome sequencing (WGS) and total RNA data derived from fresh frozen biopsies were requested from Hartwig Medical Foundation (HMF) [37]. This cohort, containing metastatic NSCLC patients, formed the discovery dataset. The external validation was extracted from Ref. [4] and consisted of whole exome sequencing (WES) of formalin-fixed paraffin-embedded (pre-immunotherapy) NSCLC samples of tumors and matched normal tissue. A more detailed description of cohort assembly can be found in the Supplementary Material.

### DNA processing

Whole genome sequencing, variant calling, and purity estimation were performed by HMF [37, 38]. Since the whole genome sequencing reads were mapped to GRCh37, we used crossmap [39] to perform a liftover (or, remapping) from GRCh37 to GRCh38 using Ensembl’s corresponding chain file.

### Mutation deconvolution

COSMIC v3.3 (June 2022) mutational signatures, ***H***, were used to deconvolute mutations (see Supplementary Table 1 for a list of analysed signatures). For substitutions and short indels, these signatures ***H*** describe the nucleotide alteration distribution [10, 40]. Release v3.3 adds the recently developed copy number signatures which capture the copy number × zygosity × length distribution [20, 21].

We first used SigProfilerMatrixGenerator on amino acid-changing mutations to extract mutation spectra of single base substitutions (singlets) with two flanking bases (SBS-96), doublet base substitutions (doublets, DB-78), and insertion deletions (indels, ID-83) [41]. The same package was used to compute genomewide copy number alterations (CN-48) [21]. Each mutation spectrum, ***X***, is a positive *m*-by-*n* matrix (i.e.,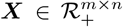) consisting of *m* samples, and *n* mutation/channels (*n* = 96, *n* = 78, *n* = 83, *n* = 48 for SBS-96, DB-78, ID-83, and CN-48, respectively) counting the number of mutations per channel. The positive mutational signature matrix ***H*** relates the *p* signatures (rows) to the corresponding *n* mutation type/channels (columns),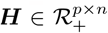. (Throughout this paper we adhere to the convention that *m, n*, and *p* indicate the number of patients, number of mutation types/channels, and number of signatures [except for *p*-values which will be clear from the context], respectively.) Using the spectrum ***X*** and mutational signatures ***H***, signature attribution ***W***, a positive *m*-by-*p* matrix (i.e.,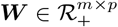) such that ***X*** ≈ ***W H***, was computed by nonnegative matrix factorisation using a coordinate descent solver with an error tolerance of 10^−6^ for no more than 10^4^ iterations.

Signature attributions that refer to possible sequencing artefacts [40] or with zero variance in either durable benefit stratum in the discovery set were excluded from the analysis. Mutational signatures of singlets, doublets, indels, and TMB (i.e., total mutation count) were obtained from non-synonymous mutations and normalised by (exome) coverage size in megabases (Mb) and rounded to two decimals to retain three significant digits. For the Hartwig WGS, the exome coverage size in individual samples was unknown. Therefore, a size of 47.9 Mb was taken [42]. For the discovery cohort (Miao *et al*. [4]), we used the size as indicated in their Supplementary Table 1. Since the copy number variants span large portions of the genome (both exonic and intronic regions), we report the total, whole genome, copy number attributions.

### RNA processing

Singlets were traced back to transcripts to study how the mutational signatures manifest at the transcription level, as a surrogate for protein expression. Out of the *m* = 101 patients in the discovery cohort, raw (total) RNA sequencing data of *m* = 40 patients were available (no RNA was available in the validation cohort). Briefly, raw sequencing data were trimmed, aligned, and converted into transcripts per million (TPM). After quality control, two inferior-quality samples and two samples with insufficient follow-up were excluded, leaving a total of *m* = 36 samples for analysis. The amount of transcripts containing a variant was re-estimated to account for differences in tumor content. RNA per signature was obtained by pooling transcripts containing the ≥ 50% dominant mutations of the given signature. A more detailed description of the method can be found in the Supplementary Information.

## Statistical Analysis

Here and in the following, all tests were two-sided.

### Univariate analysis

Differences in mutational signature distributions were determined by a Kolmogorov-

Smirnov (K-S) test. When more than one mutational signature was considered at a time, the Benjamini-Hochberg (B-H) correction was applied to control for false positive discoveries. Correlations between signatures and mutation counts per gene were evaluated using Kendall *τ* rank correlation (with B-H correction) to account for ties (both measures were derived from amino-acid sequence changing mutations only). Significantly correlated genes were subsequently annotated as (i) tumor suppressor gene using the tumor suppressor gene database website version 2.0 [43] (accessed 1st September 2022) and (ii) as oncogenes when present in the Cancer Gene Census COSMIC v.96 [44] (accessed 16th September 2022) and tested for enrichment. For differential gene expression on the transcripts with a variant, we used a non-parametric B-HK-S test because for some genes no mutated transcripts were measured (these zeros were not possible to analyse with DESeq2). We use *q* to denote the multiple testing corrected *p*-values and a significance level ≤ 5% was considered statistically significant.

### Efficacy classifier

Patients were labelled as durable benefit [progression-free survival (PFS) ≥ 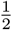 year] or non-durable benefit (PFS < 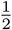 year), whenever censoring permitted unambiguous label assignment. All patients were included in PFS analysis. In view of the limited overall survival (OS) data available in the Discovery cohort, we did not consider OS as an alternative endpoint (to PFS) for our classifier. To predict DB, we used a naive Bayes classifier: a classic supervised machine learning method [45, 46] that works particularly well with few samples, even when its conditional independence assumption is violated [47]. Features were modelled with a zero-inflated exponential distribution. Results reported on the discovery cohort were obtained by leave-one-out cross-validation while inference on the validation set was done after training on the entire discovery cohort.

Estimates *a* and ninety-five per cent confidence intervals [*a* − *b, a* + *c*] of the average precision, area under the receiver operating characteristic curve (ROC AUC), *F*_1_ score, sensitivity, and specificity were estimated by bootstrapping for 1000 iterations and denoted as 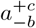. Head-to-head model comparisons were evaluated using a paired permutation test (PPT), while performance estimates of different sets were compared using an unpaired permutation test (UPT).

For the PFS analysis (including all patients), we visualised the (out-of-fold) predicted versus actual PFS outcome using the Kaplan-Meier method. To quantify agreement, hazard ratios and significance were evaluated using Cox regression (for which we tested appropriateness). To compare hazard ratios *h*_*r*_, corresponding coefficients (i.e., − ln *h*_*r*_), were compared with a regression coefficient test [48].

## Acknowledgements

We thank Fenneke Zwierenga en Benthe Muntinghe for proofreading. This publication and the underlying research are partly facilitated by Hartwig Medical Foundation and the Center for Personalized Cancer Treatment (CPCT) which have generated, analysed and made available data for this research.

## Data availability statement

The discovery data that support the findings of this study are available from Hartwig Medical Foundation (data request # 094) but restrictions apply to the availability of these data, which were used under license for the current study, and so are not publicly available. Data are however available from the corresponding author on reasonable request and with permission of Hartwig Medical Foundation.

The validation set can be extracted from Ref. [4]. Code and analysis notebooks are publicly available on Gitlab under the MIT license.

## Data ethics statement

The discovery set was collected as part of pan-cancer studies CPCT-02, DRUP and WIDE described in [37], was evaluated and approved by the medical ethical committees of University Medical Center Utrecht and the Netherlands Cancer Institute and was executed in accordance with the relevant guidelines and regulations. All data contained in the discovery set was obtained through written informed consent of all the subjects for the purpose of whole genome sequencing and data sharing for cancer research.

## Author contribution

Conceptualization: HCD, BvE, MT, GAL, HJMG; Methodology: HCD; Software: HCD; Validation: HCD; Formal analysis: HCD, BvE; Investigation: HCD; Data Curation: HCD, BvE; Writing – Original Draft: HCD; Writing – Review & Editing: HCD, BvE, MT, GAL, LCLTvK, ES, TJNH, HJMG; Visualization: HCD, BvE, TJNH; Supervision: HJMG;

