## Supporting Material for "Using genomic scars to select immunotherapy beneficiaries in advanced non-small cell lung cancer"

### A Extended Methods

#### A.1 Cohort Assembly

Patient characteristics and biopsy data were requested from Hartwig Medical Foundation (HMF), data request number 094. The HMF dataset was used as the discovery cohort and contains samples from patients with advanced cancer not curable by local treatment [1]. Patients were recruited from 41 hospitals in the Netherlands [1]. Limited survival data and no performance score, ethnic background, nor smoking status were available for analysis. Whenever available, response data (of unspecified evaluation criterion) were used to construct progression-free survival data. Eight patients without response measurement who died shortly after treatment stop were marked as progressive disease at the last follow-up. After excluding seven patients without DNA sample and one patient that was not treated with immunotherapy, we compiled a dataset with a total of $m = 101$ patients. Accrual of these patients was between January 2016 and May 2019, with the last follow-up information from May 2020. Whole DNA was extracted from freshly frozen biopsies (of unspecified histology), and total RNA isolation was performed as described previously [1].

The external validation was extracted from Ref. [2] and consisted of whole exome sequencing (WES) of formalin-fixed paraffin-embedded pre-treatment samples from immune checkpoint inhibition-treated advanced lung cancer. In turn, variants and clinical characteristics in that work consisted of newly sequenced samples combined with data from Refs. [3, 4] (all advanced lung cancer). One patient, who had small cell instead of non-small cell lung cancer, was excluded leaving $m = 56$ patients in total. No study dates were specified in Ref. [2]. No RNA was available for analysis in the validation set.

#### A.2 Upstream RNA processing

FASTQ files were processed by sample-wise pooling of reads from different lanes and subsequent trimming with Trimmomatic [5] using a four-base sliding window to cut reads falling below a quality of 25, dropping reads shorter than 50 bases, and trimming the 8 leading bases (the head) if below the quality threshold (and otherwise default parameters, as suggested in the manual). The subsequent sequencing data were analysed using FastQC and aligned with STAR [6] using the thirteenth release for the GRCh38 human reference assembly and Gencode's gene annotation, version 36, to guide read mapping. After alignment, MultiQC [7] was used to compare overall sample quality allowing us to exclude two anomalous samples of inferior quality. Two additional RNA samples were excluded from downstream analysis because the patients had insufficient follow-up to unambiguously assign a durable benefit label (leaving a total of $m = 36$). The remaining aligned samples (in coordinate sorted BAM format) were indexed using samtools [8]. The somatic DNA mutations were tracked at the RNA level by generating pile-ups at the (remapped) genomic position to obtain the variant allele frequency, $\omega_{\text{mut}}$ (un-remappable variants were discarded). The number of RNA counts were estimated per exon (as suggested in the manual) using HTSeq-count [9] with the aforementioned gene annotation. To estimate the amount of mutant RNA we first computed the number of transcripts per million, $t_\alpha$, of transcript $\alpha$. Briefly, using the RNA counts per transcript, $n^{(\alpha)}$, and the transcript length, $l^{(\alpha)}$, (in base pairs), its' value is defined by

$$t_\alpha = \frac{n^{(\alpha)}}{l^{(\alpha)}} \frac{10^6}{Z}, \tag{1}$$

with $Z$ a normalisation constant $Z = \sum_\alpha \frac{n^{(\alpha)}}{l^{(\alpha)}}$.

#### A.3 Mutated RNA re-restimation

Observe that the amount of mutated RNA molecules depends on the sample's overall tumor purity $f_t$, as well as the tendency of tumor cells to express and/or break down the mutated transcripts. We, therefore, introduce a (transcript specific) enrichment factor, $r_\alpha$, defined as the ratio between the allele frequency $\omega_{\text{mut}}^{(\alpha)}$ in transcript $\alpha$ and overall tumor purity, $r_\alpha = \frac{\omega_{\text{mut}}^{(\alpha)}}{f_t}$. This enrichment factor, $r_\alpha$, re-estimates the RNA from the samples as if they had 100% tumor purity. When a transcript contained more than one mutation, the allele frequency was, for simplicity, averaged. Finally, the estimated the amount of mutant RNA molecules is computed as:

$$n_{\text{mut}}^{(\alpha)} = r_\alpha t_\alpha. \tag{2}$$

#### A.4 RNA per signature

For each mutational signature, $i$, dominant mutations, $j$, were determined by selecting the smallest set of mutations, $T$, that account for $\geq 50\%$ of somatic mutations (i.e., $\sum_{j \in T} H_{ij} \geq 0.5$). Using these dominant mutations, RNA expression of transcripts containing mutations in set $T$ [Eq. (2)] were pooled for analysis.

#### A.5 Net benefit

For the net-benefit analysis we use the following definitions from [10] and [11]

$$\text{NB}_{\text{treated}}(t) = \frac{\text{TP}}{m} - \frac{\text{FP}}{m} \frac{t}{1-t}, \tag{3}$$

$$\text{NB}_{\text{untreated}}(t) = \frac{\text{TN}}{m} - \frac{\text{FN}}{m} \frac{1-t}{t}, \tag{4}$$

where $t$ is the probability threshold for accepting a positive prediction, and TP, FP, TN, FN are the number of true positives, false positives, true negatives, and false negatives, respectively. Here a low $t$ indicates that we attach great value to avoiding false negatives, and a high $t$ if we attach great value to avoiding false positives. The combined net benefit is defined as it's sum $\mathrm{NB}_{\mathrm{combined}}(t) = \mathrm{NB}_{\mathrm{treated}}(t) + \mathrm{NB}_{\mathrm{untreated}}(t)$. A metric that directly follows is the integrated net benefit from [12]

$$\mathrm{NBI} = \int_0^1 \mathrm{NB}(t)\mathrm{d}t\,. \tag{5}$$

A downside of the net benefit is the lack of interpretability and it does not transparently include quality-adjusted life years or financial cost. To augment the net benefit we also consider cost as a function $C(t)$ of true positives, true negatives, false positives and false negatives, assuming independent costs:

$$C(t) = \frac{\omega_{\mathrm{TN}}\mathrm{TN}(t) + \omega_{\mathrm{TP}}\mathrm{TP}(t) + \omega_{\mathrm{FN}}\mathrm{FN}(t) + \omega_{\mathrm{FP}}\mathrm{FP}(t)}{m\sum_i \omega_i}\,, \tag{6}$$

where $\boldsymbol{\omega}$ represents the weights. Here weights are taken as $\omega_{\mathrm{TN}} = 0$, $\omega_{\mathrm{TP}} = 100$, $\omega_{\mathrm{FN}} = 200$, $\omega_{\mathrm{FP}} = 100$; this can be translated as; we attach a cost of 0 to true negatives as we do not incur extra costs, a cost of 100 to an accurate estimation that immunotherapy is necessary, i.e. we attach a cost of 100 to effective immunotherapy, then a cost of 300 to a false rejection of immunotherapy to account for incurring unnecessary suffering, and again a cost of 200 for ineffective immunotherapy, the reasoning for the latter is that we have the cost of the immunotherapy plus the cost of treating the side-effects. The attribution of weights should be done with extreme care, and we want to emphasize that we merely present this as an addition to the classical net benefit analysis.

### B Extended Results

#### B.1 Correlation between signatures and prior treatment

The majority of patients in the discovery cohort had a history of chemotherapy and/or radiotherapy. Both radiation exposure [13, 14, 15] and chemotherapy [13] are known to cause distinguishing mutations. We, therefore, analysed all signatures (SBS, DB, indel, and CNV) to see if that led to any significant signature differences between the exposed versus unexposed groups. For both chemotherapy and radiotherapy, no differences between the groups was observed. Our negative findings are explained by the fact that (i) lung cancer is, after melanoma, one of the cancers with the highest mutation burden [16], (ii) signatures partially overlap, (iii) our population is small, and (iv) our false discovery control is strict. Treatment-related mutations could be buried by other mutation sources generating similar mutations — c.f., the overlap of platinum chemotherapy signature SBS35 and smoking-associated signature SBS4. The resulting mutational signature differences may therefore be too small to detect in our population, given our multiple testing correction.

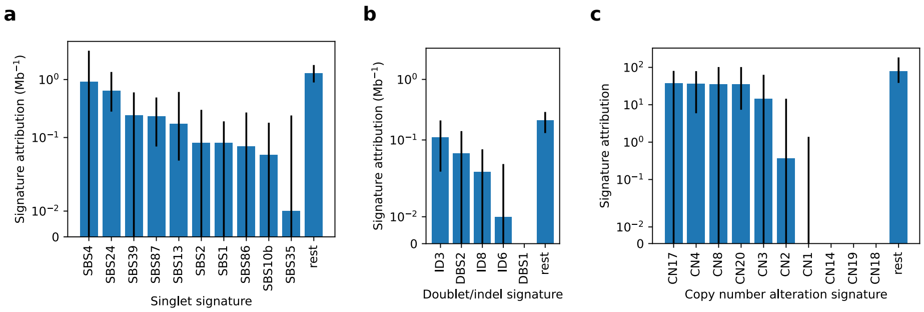


Figure S1: **Mutational signature deconvolution of tumor DNA derived from the metastatic non-small cell lung cancer discovery cohort.** **a**, Top ten non-synonymous single base substitution (singlet) signature attributions (prefix: SBS), per megabase (Mb) of exome. Rest: signature attributions of all other singlet signatures combined. **b**, Top five non-synonymous doublet base substitution (doublet) and short insert deletion (indel) signature attributions (prefixes DB and ID, respectively) per Mb of exome. Rest: signature attributions of all other doublet and indel signatures combined. **c**, Top ten whole genome copy number (CN) alteration signatures. Signatures are ranked by their total, genome wide, median attribution. Values and error bars indicate median and upper-lower quartiles, respectively.

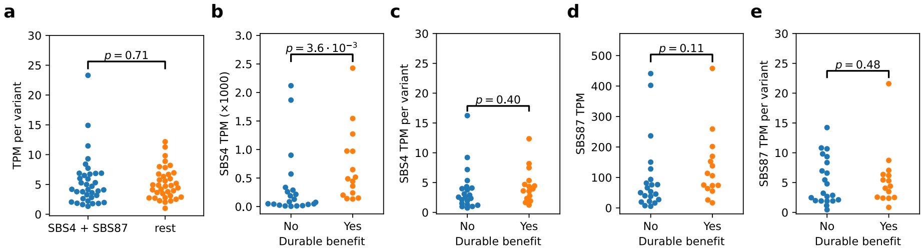


Figure S2: **Amount of mutation carrying RNA, per signature, in pre-immunotherapy tumor tissue from the discovery set.** **a**, RNA harbouring (smoking associated) signature SBS4 or (thiopurine exposure associated) signature SBS87-dominant single base substitutions (singlets) are compared to RNA from all other singlets (rest). The amount is normalised by the number of corresponding mutations. **b**, Amount of RNA harboring signature SBS4-dominant singlets split by outcome. **c**, As in **a**, but normalised by the number corresponding mutations. **d**, Amount of RNA harboring signature SBS87-dominant singlets split by outcome. **e**, As in **c**, but normalised by the number corresponding mutations. Significance was assessed using a Kolmogorov-Smirnov test. Abbreviations: TPM, transcripts per million.

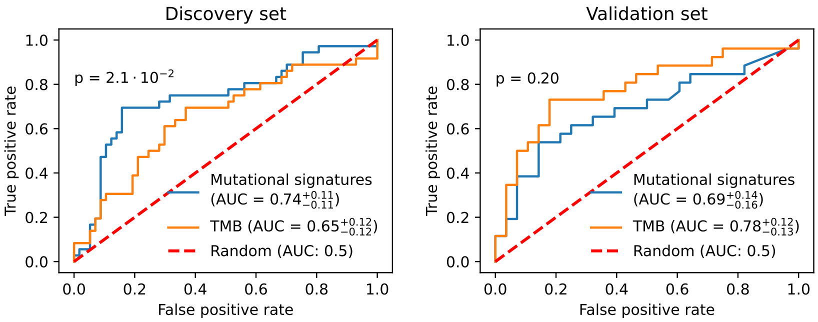


(a) Discovery dataset (Hartwig)

(b) Validation dataset [Miao *et al.* ('18)]

Figure S3: Receiver operating characteristic (ROC) curve analysis of durable benefit models that use pre-immunotherapy tumor tissue in non-small cell lung cancer. A naive Bayes classifier is trained on the discovery set using either tumor mutational burden or with the mutational signature SBS4 and SBS87-based model. In the discovery dataset (left), the ROC area under the curve (AUC) of the mutational signature-based model is significantly higher than the TMB-based model [$0.74^{+0.10}_{-0.12}$ versus $0.65^{+0.12}_{-0.13}$, respectively, $p = 0.016$ paired permutation test (PPT)]. In the validation dataset (right), there was no significant difference in the ROC AUC ($0.69^{+0.14}_{-0.14}$ versus $0.78^{+0.12}_{-0.14}$, respectively, $p = 0.18$ PPT). Estimates and corresponding 95% CIs are indicated by sub and superscripts.

#### B.2 Whole genome sequencing signatures linked to durable benefit

Signatures derived from all mutations (i.e., whole genome, including synonymous variants) found primarily smoking-related signatures, SBS4 ($q = 0.0081$, B-HK-S test), doublet base substitution DBS2 ($q = 0.0075$, B-HK-S test), and indel signature ID3 ($q = 0.044$, B-HK-S test) associated with durable benefit. While signature SBS87 was not recapitulated in WGS ($q = 1$, $p = 1.0$), we did find another doublet signature, DBS6, of unknown aetiology associated with DB ($q = 0.0090$, B-HK-S).

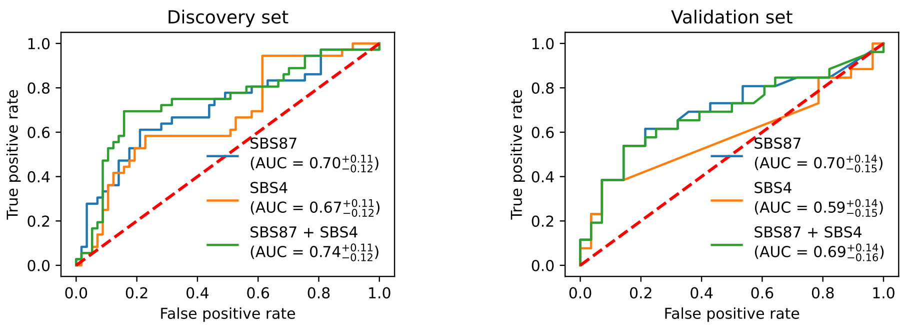


(a) Discovery dataset

(b) Validation dataset

Figure S4: Receiver operating characteristics curve analysis of naive Bayes classifier with either one mutational signature, or both. Combining both signatures slightly improves the model on the discovery set (left). Holdout predictions on the external validation (right) set shows that the SBS87 classifier performs about the same as the one combining both mutational signatures. The area under the curve and 95% confidence intervals are indicated in parentheses in the legend.

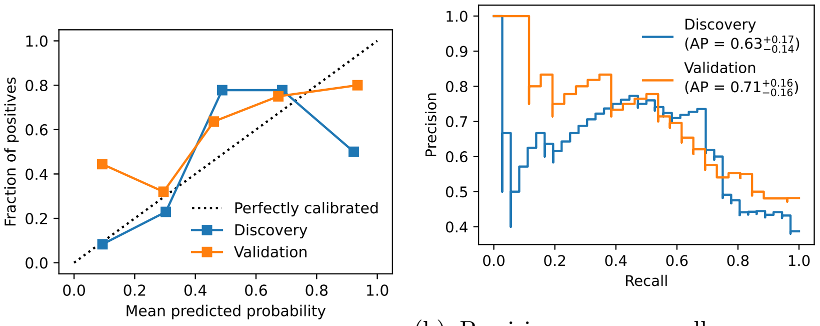


(a) Calibration curve as a function of classification probability.

(b) Precision versus recall curve parameterised by the classification probability threshold.

Figure S5: Performance of mutational signature model that predicts durable benefit from pre-immunotherapy tumor tissue in non-small cell lung cancer. Estimates and corresponding 95% confidence intervals are indicated by sub and superscripts. Abbreviations: AP, average precision.

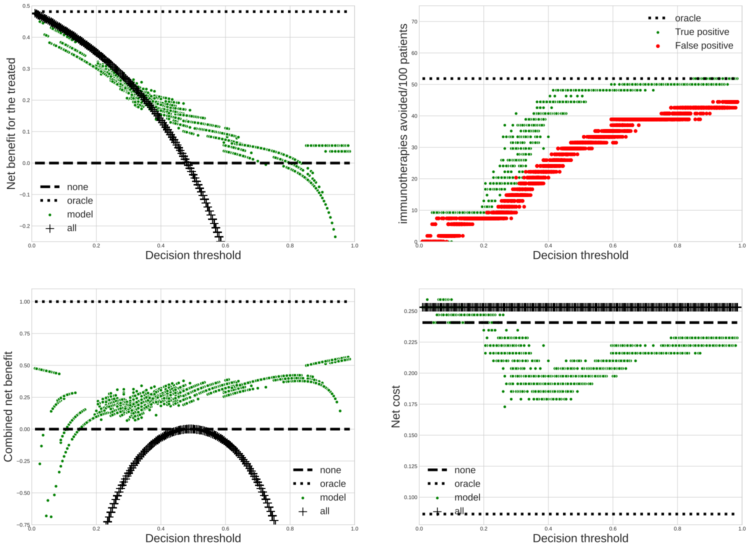


Figure S6: Net benefit plot for the discovery dataset, with three reference results, namely **none** when no patient is treated, **all** when all patients are treated and **oracle** when we have a perfect model, for the net cost curve we use $\omega_{\mathrm{TN}} = 0$, $\omega_{\mathrm{TP}} = 100$, $\omega_{\mathrm{FN}} = 300$, $\omega_{\mathrm{FP}} = 200$. The dots are obtained by resampling 50 times.

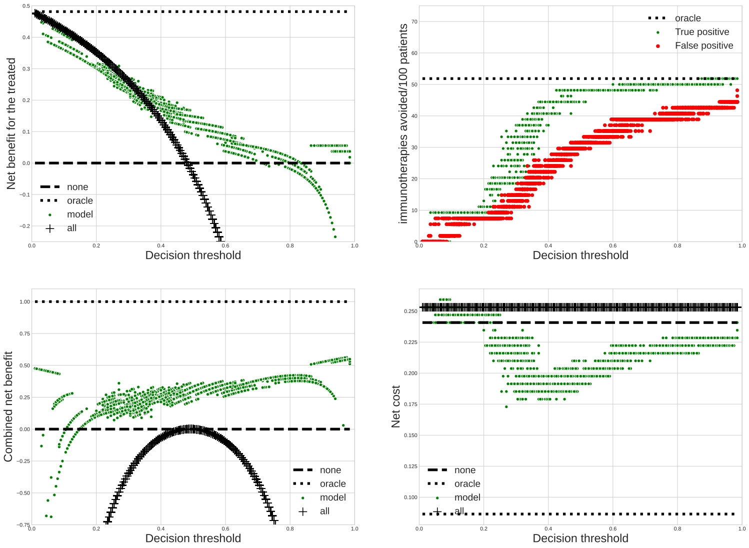


Figure S7: Net benefit plot for the validation dataset, with three reference results, namely **none** when no patient is treated, **all** when all patients are treated and **oracle** when we have a perfect model. The dots are obtained by resampling 50 times.
